# Population Genomics of the Foothill Yellow-Legged Frog (*Rana boylii*) and RADseq Parameter Choice for Large-Genome Organisms

**DOI:** 10.1101/186635

**Authors:** Evan McCartney-Melstad, Müge Gidiş, H. Bradley Shaffer

## Abstract

Genomic data are useful for attaining high resolution in population genetic studies and have become increasingly available for answering questions in biological conservation. We analyzed RADseq data for the protected foothill yellow-legged frog (*Rana boylii*) throughout its native range in California and Oregon, including many of the same localities included in an earlier study based on mitochondrial DNA. We recovered five primary clades that correspond to geographic regions within California and Oregon, with better resolution and more spatially consistent patterns than the previous study, confirming the increased resolving power of genomic approaches compared to single-locus analyses. Bayesian clustering, PCA and population differentiation with admixture analyses all indicated that approximately half the range of *R. boylii* consists of a single, relatively uniform population, while regions in the Sierra Nevada and Central Coast Range of California are deeply differentiated genetically. Additionally, a major methodological challenge for large genome organisms, including many amphibians, is deciding on sequence similarity clustering thresholds for population genetic analyses using RADseq data, and we develop a novel set of metrics that allow researchers to set a sequence similarity threshold that maximizes the separation of paralogous regions while minimizing the oversplitting of naturally occurring allelic variation within loci.

## Introduction

Amphibian population declines are a global concern, driven by habitat degradation, climate change, emerging diseases, pesticides, introduced species and pollution (Houlahan et al. 2000; Kiesecker et al. 2001; Stuart et al. 2004; Whiles et al. 2013). California has been a region of intense amphibian declines, with severe reductions across a wide range of anurans and urodeles (Hayes and Jennings 1986; Davidson 2004; Vredenburg et al. 2007; Kupferberg et al. 2012; Rogers and Peacock 2012; Ryan et al. 2014; Thomson et al. 2016).

One species that has received renewed attention from regulatory agencies is the foothill yellow-legged frog (*Rana boylii*), a species that historically was widely distributed from southwestern Oregon to southern California. This frog breeds exclusively in slow-flowing stream habitats and faces multiple threats, leading to its designation as a California Species of Special Concern (Thomson et al. 2016), its consideration for federal listing (US Fish and Wildlife Service 2015), and most recently its designation as a candidate for listing under the California Endangered Species Act in June 2017. Several *R. boylii* studies have investigated the modification and loss of its breeding habitat through the construction of dams (Lind et al. 1996, 2016; Kupferberg et al. 2012), with one set of analyses showing that the species has disappeared from at least 50% of historical localities (Davidson et al. 2002; Davidson 2004). Pesticides may be particularly problematic for this species (Davidson et al. 2002; Davidson 2004; Bradford et al. 2011; Kerby and Sih 2015), and recent studies suggest that chytridiomycosis caused by the fungus *Batrachochytrium dendrobatidis* (Bd) may constitute a significant threat to the species in some areas (Ecoclub Amphibian Group et al. 2016; Adams et al. 2017). Both experimental and field-based studies of *R. boylii* have also implicated synergistic effects of Bd infection and pesticides on metamorphic growth (Davidson et al. 2007), and of the combination of Bd infection, drought and invasive bullfrog occurrence on Bd prevalence and load (Adams *et al.* 2017), suggesting that site and lineage-specific effects may be important drivers of decline.

Given its status as a state-protected species and its consideration for federal and state listing, information regarding the spatial genetic structure of *R. boylii* is critical. A previous study using primarily mitochondrial DNA (mtDNA) found evidence of spatial genetic structure across the species, but had relatively low resolution and yielded sometimes puzzling results with respect to geography (Lind et al. 2011). However, several results from that study were broadly concordant with those from other aquatic amphibians in California, including the discovery of unique, relatively differentiated populations in the southern-most (Kern River drainage and southern Monterey County) and the northern-most (Willamette River valley in central Oregon) localities, and a very general concordance between drainage basins and genetic lineages, particularly with respect to the Sacramento-San Joaquin drainage system. However, Lind *et al.* (2011) also found that the low resolving power of a single mitochondrial marker precluded strong conclusions for conservation decision-making. We therefore applied a genomic restriction site-associated DNA sequencing (RADseq) approach to collect information for tens of thousands of nuclear loci for samples from across the range of the species, including many of the same samples and sites studied by Lind *et al.* (2011).

One liability of the RADseq approach is the anonymous nature of putative loci, and therefore the difficulty of firmly establishing homology across loci (Ilut et al. 2014; Harvey et al. 2015). Relatively few RADseq studies have been published in animals with extremely large genomes including most amphibians (but see Streicher et al. 2014; Roland et al. 2016; Nunziata et al. 2017). Although the genome size of *R. boylii* is not directly known, the closely related species *Rana pretiosa*, *Rana aurora*, and *Rana cascadae* (Macey et al. 2001; Shaffer et al. 2004) have haploid genome sizes ranging from 7.33gb to 9.13gb (Olmo 1973; Vinogradov 1998), suggesting that *R. boylii* is likely in the 7-10gb range. The choice of sequence similarity clustering threshold is central to all downstream analyses and may be particularly important in genomic RADseq studies for these and other large-genome species, as highly repetitive genomes may be susceptible to the improper clustering of paralogous regions into single loci (Ilut et al. 2014; Harvey et al. 2015). To quantitatively address this problem, we generated a set of four metrics for assessing optimal clustering thresholds in RADseq studies where large genomes may pose a concern. These metrics, which are further described in the following Methods section, were designed to optimize the threshold clustering value at which natural allelic variation in the population began to be clustered into separate genome loci, so that data assembly for population genetic analyses can utilize a value close to, but not exceeding that value.

## Materials and Methods

### Laboratory methods

Tissues from 93 *Rana boylii* samples were obtained from 50 locations (between 1 and 9 samples per locality) in California and Oregon across the geographic range of the species (Figure 1, Table 1), with priority given to samples representing each of the mtDNA clades recovered by Lind *et al.* (2011). DNA was extracted from tissues using a salt extraction protocol (Sambrook and Russell 2001). Dual-indexed sequencing libraries were prepared using the 3RAD protocol, a variant of the double-digest RADseq protocol (Peterson et al. 2012), with MspI as the common-cutting enzyme, SbfI as the rare-cutting enzyme, and ClaI as the dimer-cleaving enzyme (Hoffberg et al. 2016).

**Figure 1:**
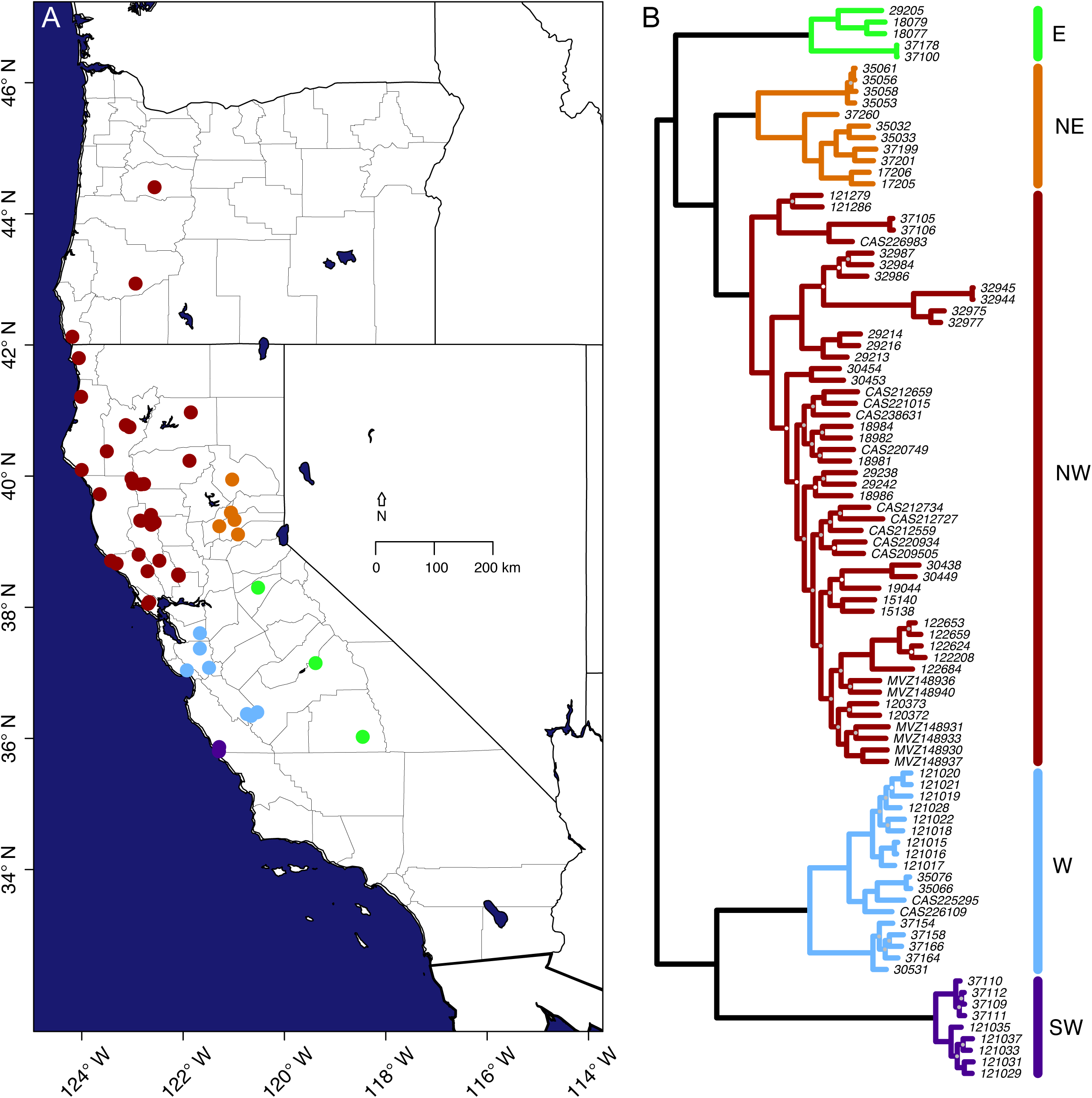
A) Map of *Rana boylii* sample localities. County outlines are shown within California and Oregon. Dots are colored by major grouping as determined in the B) RAxML phylogeny, with midpoint rooting. Nodes without dots received 100% bootstrap support, those with white dots received less than 95% but greater than or equal to 75% bootstrap support, while nodes with grey dots received less than 75% bootstrap support.

**Table 1:** Sample information. Sample names with the prefix HBS indicate samples from the Shaffer Lab collection at UCLA, MVZ denotes samples from the Museum of Vertebrate Zoology, University of California, Berkeley, and CAS denotes samples from the California Academy of Sciences.

| Sample | Latitude | Longitude | TreeMix ID | State | County | Basin |
| --- | --- | --- | --- | --- | --- | --- |
| HBS120372 | 38.802 | -122.879 | locality_1 | CA | Sonoma | 180101 |
| HBS120373 | 38.802 | -122.879 | locality_1 | CA | Sonoma | 180101 |
| HBS121015 | 37.037 | -121.927 | locality_2 | CA | Santa Cruz | 180600 |
| HBS121016 | 37.037 | -121.927 | locality_2 | CA | Santa Cruz | 180600 |
| HBS121017 | 37.037 | -121.927 | locality_2 | CA | Santa Cruz | 180600 |
| HBS121018 | 37.037 | -121.927 | locality_2 | CA | Santa Cruz | 180600 |
| HBS121019 | 37.037 | -121.927 | locality_2 | CA | Santa Cruz | 180600 |
| HBS121020 | 37.037 | -121.927 | locality_2 | CA | Santa Cruz | 180600 |
| HBS121021 | 37.037 | -121.927 | locality_2 | CA | Santa Cruz | 180600 |
| HBS121022 | 37.037 | -121.927 | locality_2 | CA | Santa Cruz | 180600 |
| HBS121028 | 37.037 | -121.927 | locality_2 | CA | Santa Cruz | 180600 |
| HBS121029 | 35.867 | -121.289 | locality_3 | CA | Monterey | 180600 |
| HBS121031 | 35.867 | -121.289 | locality_3 | CA | Monterey | 180600 |
| HBS121033 | 35.867 | -121.289 | locality_3 | CA | Monterey | 180600 |
| HBS121035 | 35.867 | -121.289 | locality_3 | CA | Monterey | 180600 |
| HBS121037 | 35.867 | -121.289 | locality_3 | CA | Monterey | 180600 |
| HBS121279 | 40.777 | -123.132 | locality_4 | CA | Trinity | 180102 |
| HBS121286 | 40.747 | -123.061 | locality_5 | CA | Trinity | 180102 |
| HBS122208 | 38.490 | -122.091 | locality_6 | CA | Solano | 180201 |
| HBS122624 | 38.479 | -122.087 | locality_7 | CA | Solano | 180201 |
| HBS122653 | 38.495 | -122.101 | locality_8 | CA | Solano | 180201 |
| HBS122659 | 38.499 | -122.100 | locality_9 | CA | Solano | 180201 |
| HBS122684 | 38.486 | -122.087 | locality_10 | CA | Solano | 180201 |
| HBS15138 | 38.708 | -123.423 | locality_11 | CA | Sonoma | 180101 |
| HBS15140 | 38.665 | -123.314 | locality_12 | CA | Sonoma | 180101 |
| HBS17205 | 39.109 | -120.918 | locality_13 | CA | Placer | 180201 |
| HBS17206 | 39.109 | -120.918 | locality_13 | CA | Placer | 180201 |
| HBS18077 | 38.298 | -120.522 | locality_14 | CA | Calaveras | 180400 |
| HBS18079 | 38.298 | -120.522 | locality_14 | CA | Calaveras | 180400 |
| HBS18981 | 40.377 | -123.507 | locality_15 | CA | Trinity | 180101 |
| HBS18982 | 40.377 | -123.507 | locality_15 | CA | Trinity | 180101 |
| HBS18984 | 40.377 | -123.507 | locality_15 | CA | Trinity | 180101 |
| HBS18986 | 39.723 | -123.647 | locality_16 | CA | Mendocino | 180101 |
| HBS19044 | 38.549 | -122.703 | locality_17 | CA | Sonoma | 180101 |
| HBS29205 | 37.147 | -119.387 | locality_18 | CA | Fresno | 180400 |
| HBS29213 | 41.799 | -124.059 | locality_19 | CA | del Norte | 180101 |
| HBS29214 | 41.799 | -124.059 | locality_19 | CA | del Norte | 180101 |
| HBS29216 | 41.799 | -124.059 | locality_19 | CA | del Norte | 180101 |
| HBS29238 | 40.093 | -124.003 | locality_20 | CA | Humboldt | 180101 |
| HBS29242 | 40.093 | -124.003 | locality_20 | CA | Humboldt | 180101 |
| HBS30438 | 38.052 | -122.692 | locality_21 | CA | Marin | 180500 |
| HBS30449 | 38.078 | -122.670 | locality_22 | CA | Marin | 180500 |
| HBS30453 | 41.209 | -124.014 | locality_23 | CA | Humboldt | 180101 |
| HBS30454 | 41.209 | -124.014 | locality_23 | CA | Humboldt | 180101 |
| HBS30531 | 36.398 | -120.538 | locality_24 | CA | Fresno | 180300 |
| HBS32944 | 44.405 | -122.565 | locality_25 | OR | Linn | 170900 |
| HBS32945 | 44.405 | -122.565 | locality_25 | OR | Linn | 170900 |
| HBS32975 | 42.935 | -122.939 | locality_26 | OR | Douglas | 171003 |
| HBS32977 | 42.935 | -122.939 | locality_26 | OR | Douglas | 171003 |
| HBS32984 | 42.125 | -124.187 | locality_27 | OR | Curry | 171003 |
| HBS32986 | 42.125 | -124.187 | locality_27 | OR | Curry | 171003 |
| HBS32987 | 42.125 | -124.187 | locality_27 | OR | Curry | 171003 |
| HBS35032 | 39.443 | -121.058 | locality_28 | CA | Yuba | 180201 |
| HBS35033 | 39.443 | -121.058 | locality_28 | CA | Yuba | 180201 |
| HBS35053 | 39.947 | -121.033 | locality_29 | CA | Plumas | 180201 |
| HBS35056 | 39.947 | -121.033 | locality_29 | CA | Plumas | 180201 |
| HBS35058 | 39.947 | -121.033 | locality 29 | CA | Plumas | 180201 |
| HBS35061 | 39.947 | -121.033 | locality 29 | CA | Plumas | 180201 |
| HBS35066 | 37.603 | -121.670 | locality 30 | CA | Alameda | 180500 |
| HBS35076 | 37.603 | -121.670 | locality 30 | CA | Alameda | 180500 |
| HBS37100 | 36.023 | -118.457 | locality 31 | CA | Tulare | 180300 |
| HBS37105 | 40.971 | -121.849 | locality 32 | CA | Shasta | 180200 |
| HBS37106 | 40.971 | -121.849 | locality 32 | CA | Shasta | 180200 |
| HBS37109 | 35.802 | -121.294 | locality 33 | CA | Monterey | 180600 |
| HBS37110 | 35.802 | -121.294 | locality 33 | CA | Monterey | 180600 |
| HBS37111 | 35.802 | -121.294 | locality 33 | CA | Monterey | 180600 |
| HBS37112 | 35.802 | -121.294 | locality 33 | CA | Monterey | 180600 |
| HBS37154 | 36.343 | -120.659 | locality 34 | CA | San Benito | 180600 |
| HBS37158 | 36.343 | -120.659 | locality 34 | CA | San Benito | 180600 |
| HBS37164 | 36.371 | -120.741 | locality 35 | CA | San Benito | 180600 |
| HBS37166 | 36.371 | -120.741 | locality 35 | CA | San Benito | 180600 |
| HBS37178 | 36.023 | -118.457 | locality 31 | CA | Tulare | 180300 |
| HBS37199 | 39.331 | -120.986 | locality 36 | CA | Nevada | 180201 |
| HBS37201 | 39.331 | -120.986 | locality 36 | CA | Nevada | 180201 |
| HBS37260 | 39.235 | -121.287 | locality 37 | CA | Yuba | 180201 |
| CAS209505 | 39.320 | -122.847 | locality 38 | CA | Lake | 180101 |
| CAS212559 | 39.407 | -122.635 | locality 39 | CA | Glenn | 180201 |
| CAS212659 | 39.873 | -122.832 | locality 40 | CA | Tehama | 180201 |
| CAS212727 | 39.258 | -122.628 | locality 41 | CA | Colusa | 180201 |
| CAS212734 | 39.291 | -122.562 | locality 42 | CA | Colusa | 180201 |
| CAS220749 | 39.963 | -123.020 | locality 43 | CA | Mendocino | 180101 |
| CAS220934 | 39.318 | -122.834 | locality 44 | CA | Lake | 180101 |
| CAS221015 | 39.879 | -122.773 | locality 45 | CA | Tehama | 180201 |
| CAS225295 | 37.370 | -121.671 | locality 46 | CA | Santa Clara | 180500 |
| CAS226109 | 37.077 | -121.490 | locality 47 | CA | Santa Clara | 180500 |
| CAS226983 | 40.233 | -121.876 | locality 48 | CA | Tehama | 180201 |
| CAS238631 | 39.887 | -122.988 | locality 49 | CA | Mendocino | 180101 |
| MVZ:Herp:148930 | 38.709 | -122.470 | locality 50 | CA | Lake | 180201 |
| MVZ:Herp:148931 | 38.709 | -122.470 | locality 50 | CA | Lake | 180201 |
| MVZ:Herp:148933 | 38.709 | -122.470 | locality 50 | CA | Lake | 180201 |
| MVZ:Herp:148936 | 38.709 | -122.470 | locality 50 | CA | Lake | 180201 |
| MVZ:Herp:148937 | 38.709 | -122.470 | locality 50 | CA | Lake | 180201 |
| MVZ:Herp:148940 | 38.709 | -122.470 | locality 50 | CA | Lake | 180201 |

Individual libraries were pooled at equimolar ratios and size selected in a single batch to between 500 and 700 bp using a Pippin Prep (Sage Science, Beverly, MA). Libraries were sequenced on two 100bp PE HiSeq 2000 lanes. However, the R2 failed on one of these lanes, so we retained only the R1s from both lanes to take advantage of the increased depth from two lanes of sequencing. For both lanes, custom sequencing primers were used that overlapped the restriction sites so that the first base of each read was the first base following the restriction site.

### Clustering thresholds

Previous work in *Rana boylii* showed that intron 5 of the tropomyosin gene contained only 2 variable sites across 517 bp (0.39% divergence) for a similar geographic sampling (Lind et al. 2011), which might suggest a decreased risk of oversplitting loci at higher clustering thresholds, while the larger genome sizes in amphibians with their associated increase of paralogous regions (Smith et al. 2009) suggests an increased risk of undersplitting at lower clustering thresholds. The software pyRAD filters out paralogs using multiple approaches, including flagging loci with more than two haplotypes (biologically impossible for a single locus in a diploid organism) and identifying loci with a higher than expected number of heterozygous sites per individual (combined paralogous loci will appear heterozygous at sites with fixed differences between those loci). While approaches such as these have shown to be effective in other species (Ilut et al. 2014), their efficacy depends on the relative age of paralog duplication events (and therefore the amount of divergence between duplicate copies) compared to the levels of natural single-copy genetic variation present in the system, and their efficacy for distinguishing paralogs in large-genome amphibians has not been evaluated.

Although the genome size of *Rana boylii* has not been characterized, the closely related species *R. pretiosa, R. aurora*, and *R. cascadae* have been measured at between 7.33 and 9.13 billion bases (Sexsmith 1968; Olmo 1973; Vinogradov 1998; Macey et al. 2001; Shaffer et al. 2004; Gregory 2016), which is several times larger than most vertebrates. Given that such large genomes often reflect ancient duplications, we paid special attention to the possibility of improperly combining duplicated genomic regions into single clusters in our analysis. Prior research in this area has largely focused on optimizing clustering thresholds, recognizing that lower clustering thresholds potentially combine more divergent paralogs while higher thresholds correctly split them apart, potentially at the cost of splitting single-copy loci that contain highly divergent alleles (Ilut et al. 2014; Harvey et al. 2015). One favored approach is to choose the level at which the number of recovered loci or heterozygosity roughly plateaus (Ilut et al. 2014), and we employed this here (Figure 2A). We also calculated the fraction of loci flagged by pyRAD as being potential paralogs through its ploidy and locus heterozygosity filters, as this gives a sense of the fraction of loci that have been improperly lumped (and therefore the distribution of within-individual paralog genetic distances), with the understanding that these filters will not catch all true biological paralogs.

**Figure 2:**
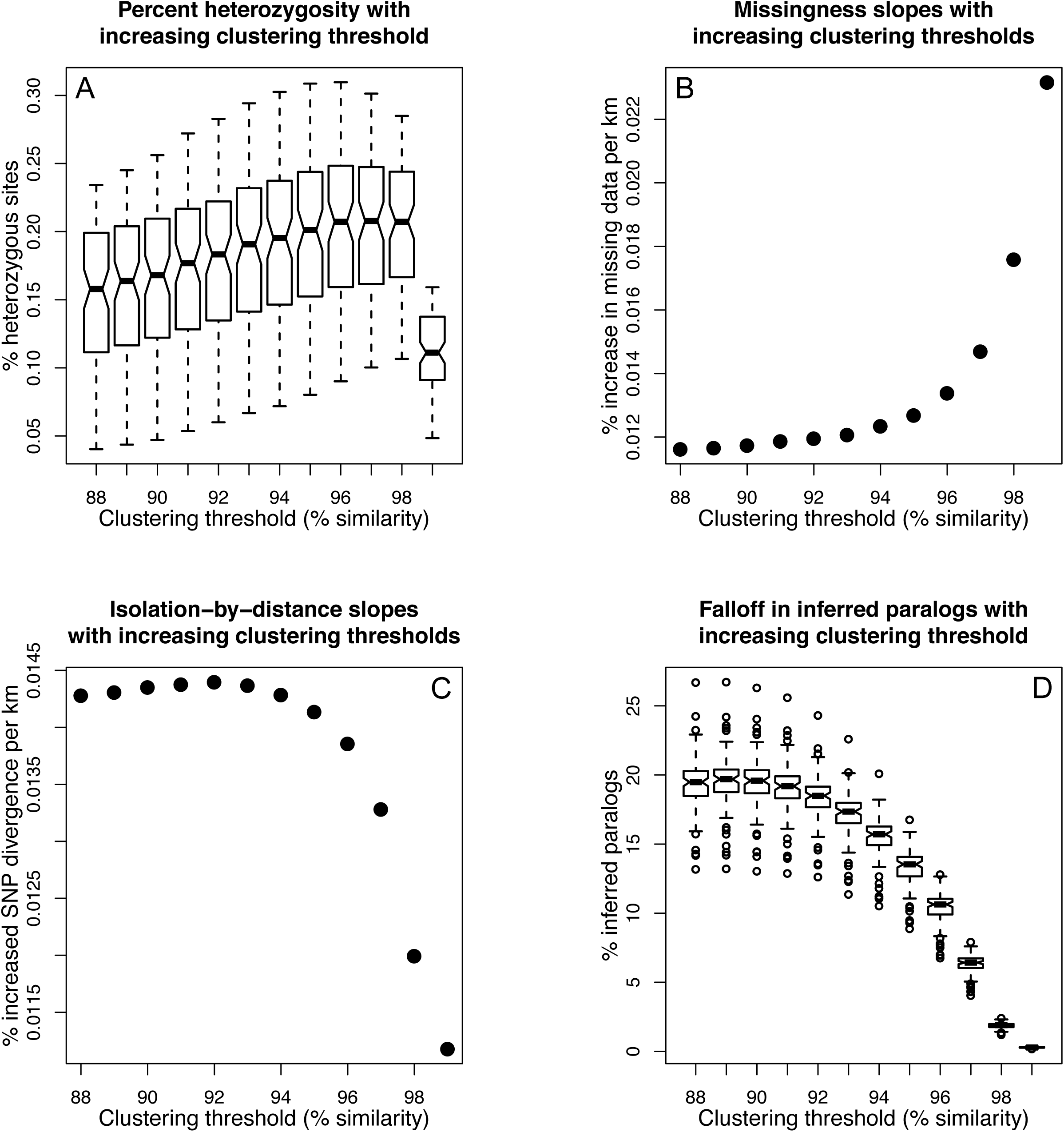
Evaluation of pyRAD clustering thresholds. A) Heterozygosity values, B) slopes of pairwise genetic data missingness between samples as a function of geographic distance, C) slopes of genetic isolation by geographic distance across all 93 samples, and D) percentage of loci flagged and removed by pyRAD as paralogs at different clustering thresholds. The notched box whisker plots showing heterozygosity and the percentage of inferred paralogs represent the individual values of each of 93 samples. Box boundaries represent the 25% to 75% interval, while the notched region represents a 95% confidence interval around the median of each group of 93 sample observations. Whiskers represent the most extreme values within 1.5 times the interquartile range from the upper and lower quartiles, and dots represent outlier values more extreme than 1.5 times the interquartile range.

Oversplitting loci with highly divergent alleles can mask true divergence by designating allelic variants of the same locus as non-homologous loci (Leaché et al. 2015). One way to test if this is a problem in a dataset is to evaluate biases in missing data as clustering thresholds increase along a suspected main axis of variation (i.e. between geographically distant samples). If, for instance, a locus is 4% divergent between two samples at opposite ends of the geographic range, then the locus will be improperly split into two distinct loci if a clustering threshold of 97% is used, with each sample containing data for one of the two loci and missing data for the other. Another component of “data missingness” between samples is mutations occurring within restriction sites, the probability of which should, on average, increase with sequence divergence. This is a biological phenomenon that should impart a positive slope to missingness plotted against geographic distance (assuming an isolation by distance model) and is not affected by clustering threshold. The increase of this slope as clustering thresholds increase, therefore, may be considered the result of oversplitting the most divergent loci amongst the most divergent (genetically and, often, geographically) samples. We also evaluated the slopes of genetic isolation by geographic distance across the different clustering thresholds. Again, we reasoned that the most geographically distant samples would be most likely to contain the greatest sequence divergence, and thus pairs of individuals that are further away from one another would be more likely to have their respective allelic variants falsely split into separate clusters, leading to artificially suppressed genetic distance calculations and a flattening of the slope of genetic isolation by distance. Thus, we tested to see at what clustering threshold the relationship between geographic distance and genetic distance became less steep, and the threshold at which pairwise data missingness drastically increased. A similar approach would be to utilize the parameters that maximize the distance/Fst ratio among the major groups under study (Ilut et al. 2014; Rodríguez-Ezpeleta et al. 2016), although this requires *a priori* assignment of genetic groups. Taken together, these factors give a sense of the clustering threshold at which paralogous loci stop being erroneously combined (undersplitting) and true single-copy allelic variants are improperly separated into non-homologous clusters (oversplitting). To implement these approaches, sequence reads were quality filtered, clustered into loci, and genotyped using pyRAD v3.0.66 (Eaton 2014). We evaluated clustering thresholds between 88% and 99% using the four above metrics by parsing the pyRAD statistics and output files.

### Quality control and clustering

To generate the final genetic dataset used for population genetic and phylogeographic analyses, pyRAD was run with the following parameters: #8/Mindepth=10, #10/Wclust=0.96, #11/Datatype=ddrad, #12/MinCov=4, #13/MaxSH=55, #24/MaxH=2. These parameters correspond to a minimum depth of 10 to include a locus for a sample, a clustering threshold of 96% similarity, a requirement that at least four samples have data to retain a locus, a maximum number of individuals being heterozygous for a particular base of 55, and a requirement that samples contain no more than two heterozygous sites at a locus to retain that locus. Mindepth and MinCov are filters for data quality and missingness, respectively, while MaxSH and MaxH serve as filters against potential paralogs being combined in a cluster.

### Data analysis

Phylogenetic analysis was conducted on a concatenated sequence alignment with a maximum of 50% missing data using RAxML v8.2.8 (Stamatakis 2014). The best-scoring maximum likelihood (ML) tree was found using 20 ML searches with the GTRGAMMA model, and 100 rapid bootstrap searches were conducted to assess node confidence.

Population structure was evaluated using fastStructure v1.0 (Raj et al. 2014). First, loci were retained that contained up to 80% missing data, as tests of population differentiation have been shown to perform relatively well at this level (Fu 2014). Then, to reduce the effects of physical linkage among variants, a single SNP was randomly chosen from each remaining RAD locus, leaving 38,520 biallelic SNPs for fastStructure analyses across all samples. Ten different random number seeds were used for K values between 1 and 12 to determine the most likely value for K to explain population structure among all 93 samples as determined by marginal likelihood. Population structure analyses often return the deepest hierarchical structure present in the data and ignore finer scale structure within the recovered clusters (Janes et al. in press; Vähä et al. 2007). To ensure that these finer, more subtle levels of population structure were identified, samples were divided into groups based upon their initial fastStructure population assignment, fastStructure was rerun for each initial group for 10 seeds ranging from K=1 to K=12, and this process was repeated recursively for additionally discovered groups, stopping when any K value with the greatest marginal likelihood was K=1 or equal to the number of sampling localities in the sample subset.

PCA was performed using SNPRelate v1.6.4 (Zheng et al. 2012) on the same set of 38,520 biallelic SNPs with a single SNP per RAD locus and no more than 80% missing data. Principal components were plotted against one another to visualize patterns of genetic variation using R 3.4 (R Core Team 2017).

We used TreeMix v1.13 (Pickrell and Pritchard 2012) to model genetic drift among sample localities while explicitly accounting for admixture. TreeMix was run with 10 different random number seeds for between 0 and 8 added migration edges. Individuals from the same locality were pooled together into 50 “population” samples, and sample size correction was turned off, as 28 of the localities consisted of a single individual. SNPs were included in the TreeMix analysis if at least one sample from each locality had data (6,004 SNPs total). Locality topologies and admixture arrows were compared between random number seeds, and random seeds with the highest likelihood for each number of migration edges are presented. The three-population test (Reich et al. 2009) as implemented in TreeMix v1.13 was also used to test for gene flow amongst the five major groups identified by RAxML and PCA using the 26,240 SNPs that contained data for at least one individual per group.

Following Lind et al. (2011), AMOVA was used to assess the influence of hydrological boundaries in structuring *R. boylii* populations (Excoffier et al. 1992). Samples were grouped according to the Watershed Boundary Database (USDA-NRCS et al. 2016) into drainage basins (6-digit hydrological unit codes, Table 1). AMOVA was conducted in Arlequin 3.5.2.2 (Excoffier and Lischer 2010) with the following hierarchical levels: 1) basin, locality, individual, and 2) five major phylogenetic clade membership, locality, individual. Individual pairwise sequence divergences were calculated using dnadist from Phylip v3.696 with JC69 distances (Jukes and Cantor 1969; Felsenstein 1989), and among-group divergences were calculated by averaging all of the individual-individual pairwise distances between group members. We also estimated Fst between the five major clades recovered by RAxML and between all sampling localities using the Weir and Cockerham (1984) method in SNPRelate 1.6.4. Tajima’s D and nucleotide diversity (π) were calculated as averages across the set of RAD loci not missing any data in each of the five major RAxML clades using VCFtools v0.1.15 (Danecek et al. 2011).

## Results

A total of 394,983,404 single-end 100bp sequence reads were used for this experiment that were derived from 93 samples across two HiSeq 2000 lanes. Samples received between 2,353,591 and 6,263,131 reads each (mean=4,247,133, stdev=840,333). Reads with more than four low-quality bases (phred scores below 20) were discarded, which reduced our data set to an average of 3,056,113 single-end reads per frog for analysis (min=1,698,025, max=4,560,049, SD=606,497).

### Clustering threshold levels

Results from pyRAD runs at clustering thresholds from 0.88 to 0.99 showed a pattern of steadily increasing heterozygosity until 0.96, where it leveled off until a sharp decrease at 0.99 (Figure 2A). Similarly, pairwise data missingness tended to increase more sharply with increasing geographic distance starting at 0.96 (Figure 2B), and the slope of genetic isolation by geographic distance began to drop precipitously at about the same level (Figure 2C). Additionally, the correlation of pairwise missingness with geographic distance increased with the clustering threshold, while the correlation of genetic distance with geographic distance increased from 88% to 90%, stayed level until 92%, decreased slightly until 97%, then fell precipitously to 99% (Figure S1). Finally, the percentage of loci flagged as paralogous based on more than two haplotypes in a diploid individual was reduced by roughly half at 0.96 compared to 0.88 (Figure 2D). Across all of the measured values (Figure 2A-D), a clustering threshold of approximately 96% or 97% represented a turning point after which our metrics deteriorated. We chose the conservatively lower value of 96% to generate the final dataset used for population genetic analyses (Ilut et al. 2014; Harvey et al. 2015).

After processing with a clustering threshold of 0.96 in pyRAD, 106,000 loci were recovered that contained data for at least four samples (Table 2). Because missing data among individuals is often high in RADseq studies (Eaton 2014) including ours, we generated two subsets of our data. The first, for RAxML analysis, contained no more than 50% missing data for any locus, resulting in 25,569 loci (2,435,662bp per individual, Table 2). The second, for all other analyses, had no more than 80% missing data across all samples, and consisted of 38,575 loci with at least one SNP.

**Table 2:** Sequence clustering and polymorphic site totals

|  |  |
| --- | --- |
| Total qualifying RAD loci (≥ four samples represented) | 106,000 |
| Total bp in RAXML sequence alignment (maximum 50% missing data) | 2,435,662 |
| Num. RAD loci with no missing data | 4,989 |
| Num. RAD loci with $< 10\%$ missing data | 14,444 |
| Num. RAD loci with $< 50\%$ missing data | 25,565 |
| Total variable sites | 207,343 |
| Total parsimony-informative sites | 128,543 |
| Sampled unlinked biallelic SNPs | 74,545 |

### RAxML

We highlight five main monophyletic groups identified in the RAxML tree that each have 100% bootstrap support (Figure 1B) and are geographically cohesive (Figure 1A). The deepest split in this tree is between coastal populations south of San Francisco Bay (the blue + purple clade in Figure 1B) and the rest of the species’ range. Within the first of these clades, two reciprocally monophyletic groups stand out. The first (purple in Figure 1; the “southwestern California” clade) is from two localities in the Central Coast Range of California in Monterey County. The second group (blue in Figure 1; the “western California” clade) consists of seven localities from Alameda, Santa Clara, Santa Cruz, San Benito and western Fresno Counties in central California. It is bounded by the Salinas River Valley to the south and west, and San Francisco Bay to the north, and occurs in both coastal and interior flowing drainages.

This inclusive group is sister to a clade consisting of all populations north of San Francisco Bay plus all Sierran *R. boylii* populations, and consists of three primary subgroups. The first (green in Figure 1; the “eastern California” clade) consists of three widely spaced localities in west-flowing drainages on the east side of California’s Central Valley in Tulare, Calaveras, and eastern Fresno Counties. The second group (red in Figure 1; the “northwestern California/Oregon” clade) consists of 33 localities extending from north of San Francisco Bay through western and central California into Oregon (including Marin, Sonoma, Solano, Lake, Colusa, Glenn, Mendocino, Tehama, Humboldt, Trinity, Shasta, and Del Norte Counties in California, and Curry, Douglas, and Linn Counties in Oregon). The final group (orange in Figure 1; the “northeastern California” clade) consists of five localities located between the eastern margins of the northwestern California/Oregon clade and the northern margin of the eastern California clade in Nevada, Placer, Yuba, and Plumas Counties in California’s Sierra Nevada.

RAxML recovered the southwestern California clade as sister to the western California clade, with the longest branch between these samples and all others. The eastern California clade was sister to a clade consisting of the reciprocally monophyletic northwestern California/Oregon and northeastern California clades (Figure 1B). The relationships among these groups were fully supported (bootstrap values of 100), but many of the shallower nodes within the five main clades were not as well supported. Bootstrap support values within the different groups differed: 0 of 3, 1 of 9, 4 of 7, 7 of 16, and 29 of 48 nodes received less than 95% bootstrap support in the eastern California, northeastern California, southwestern California, western California, and northwestern California/Oregon clades, respectively (grey and white nodes in Figure 1B).

### PCA and fastStructure

Principal components analysis broadly supported the phylogenetic results from RAxML. PC1 and PC2 (explaining 11.3% and 8.7% of the variance, respectively) separated the southwestern California and western California clades from each other and all other samples (Figure 3A). PC3 (7.1% of the variance) separated the eastern California samples from all others, while PC4 (3.9% of the variance) distinguished the northeastern California samples from the northwestern California/Oregon and eastern California samples (Figure 3B). PCs 5 through 8 (3.2% to 2.4% of the variance) suggest further substructure within the five main clades, as, for instance, PC5 is a major axis of genetic variation within the northwest California samples, separating the localities in Oregon from those further south (Figure 3C and D). Similarly, PCs 6 and 7 break the western California and eastern California samples into two groups, respectively.

**Figure 3:**
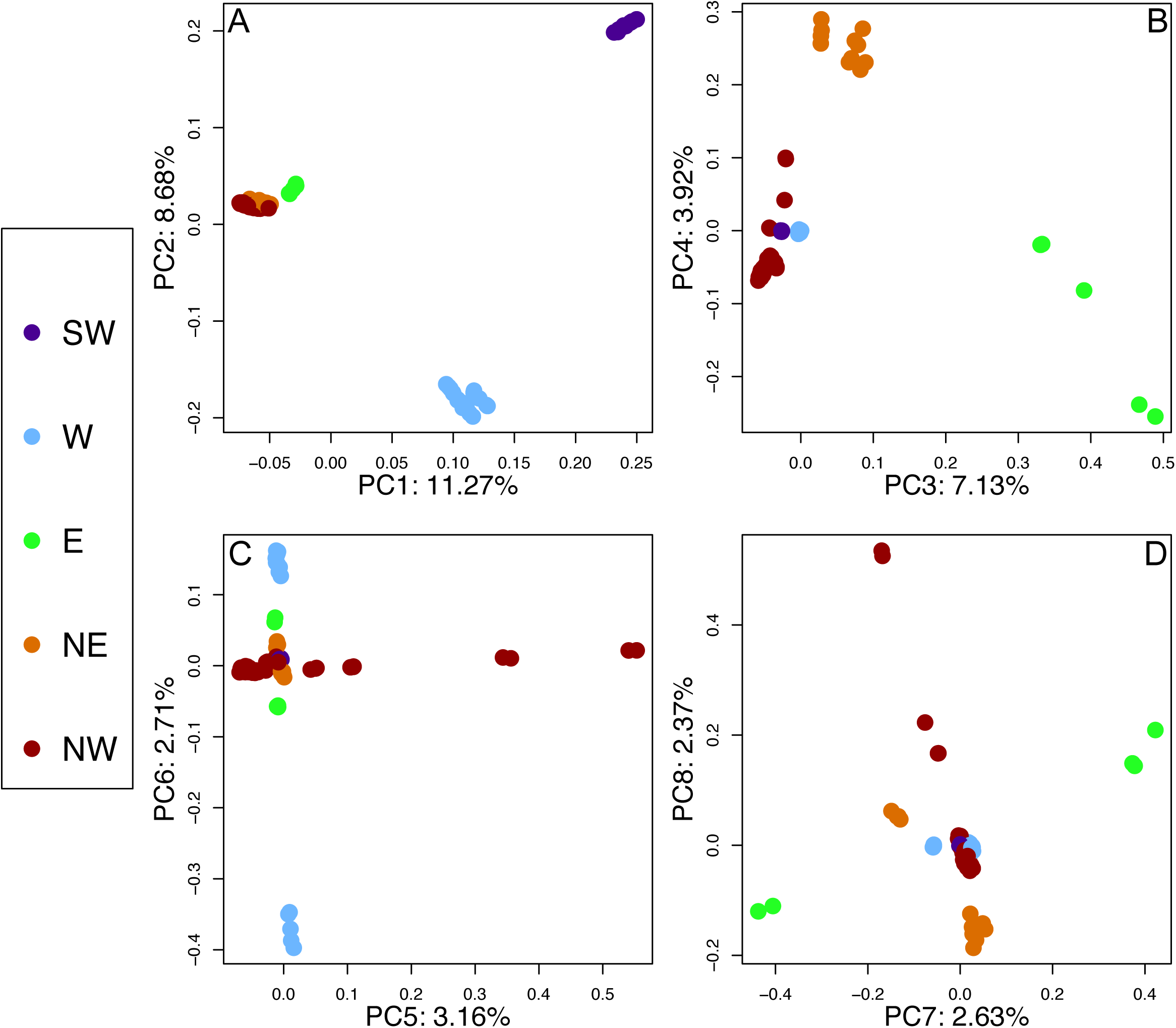
Principal components analysis. SW=southwestern California clade, W=western California clade, E=eastern California clade, NE=northeastern California clade, NW=northwestern California/Oregon clade.

Within-clade substructure was further characterized by hierarchical fastStructure analysis and was broadly concordant with the RAxML tree. For each of the ten random seeds tested, K=4 was the configuration that maximized the marginal likelihood and was also the number of model components used to explain structure in the data (Figure 4). The western California and southwestern California clades formed their own respective fastStructure populations and showed no evidence of admixed individuals. The remaining 66 individuals clustered into two populations, with 50 individuals typically clustering as purely northwest California/Oregon-clade animals and 16 individuals forming the fourth cluster, consisting of all individuals derived from the eastern + northeastern RAxMl clades. This latter group sometimes showed slight admixture from the northwest California/Oregon clade (Figure 4-1). The individuals that exhibited detectable admixture among the different random number seeds belong to the northeastern California clade, occurring in Yuba, Placer, Nevada, and Plumas Counties in California. They also all belong to a single clade representing the sister group to the northwestern California/Oregon samples in the RAxML tree (Figure 1B).

**Figure 4:**
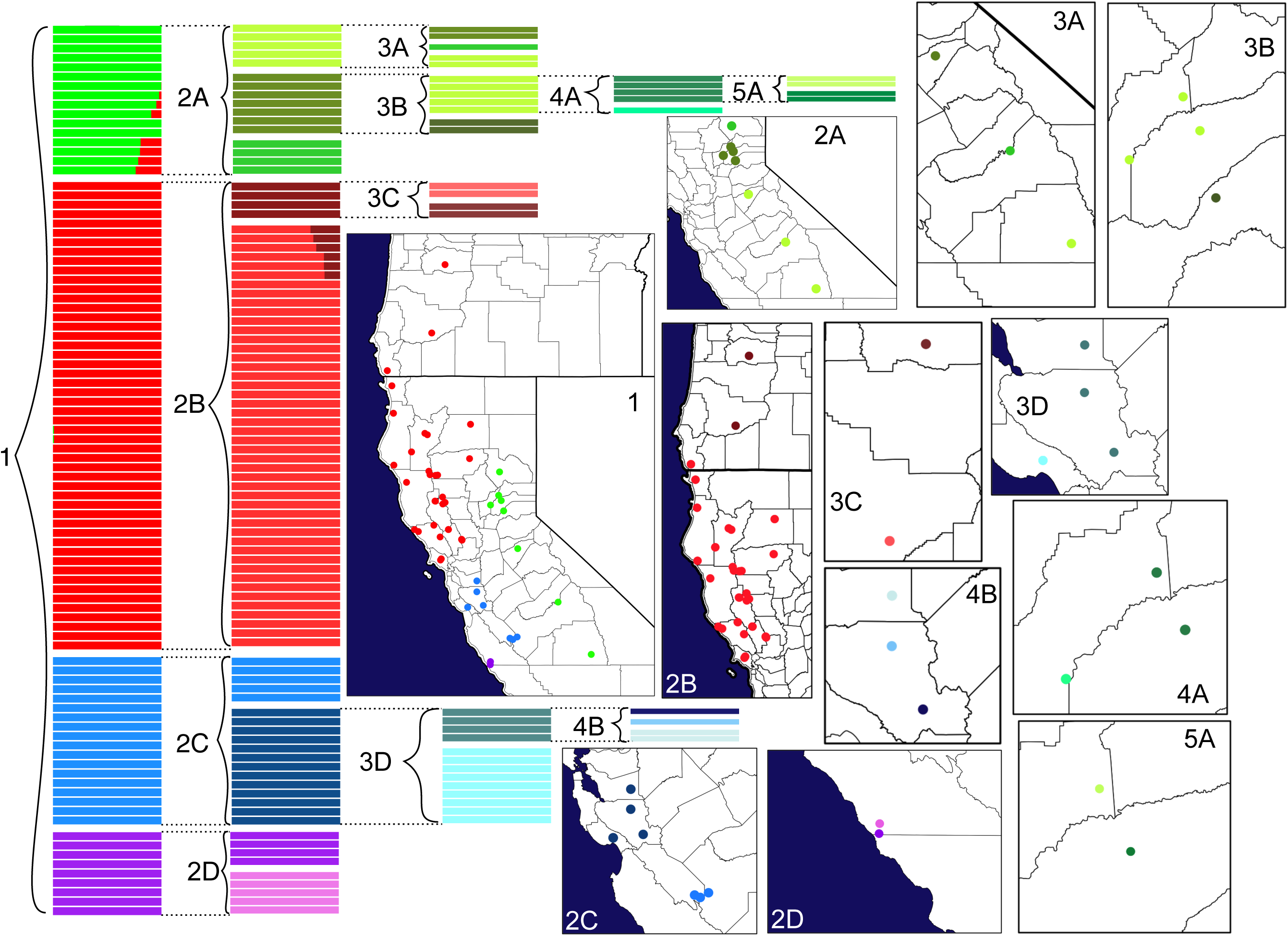
Hierarchical fastStructure results. Each number/letter combination represents a single fastStructure run and has a corresponding map with matching colors representing the samples in the fastStructure plot.

Major groups identified from fastStructure runs were then isolated and recursively subjected to fastStructure analyses until the highest marginal likelihoods were observed for K values equal to one or the number of sampling localities. The four groups identified by the first, global fastStructure analysis differed greatly in their individual substructure (Figure 4, Table S1). For instance, the global fastStructure population consisting of the eastern California and northeastern California clades required four additional rounds of fastStructure to reach the stopping condition, yielding eight distinguishable groups (Table 5). The population corresponding to the northwestern California/Oregon samples, however, required only two iterations to reach the stopping conditions and yielded just three distinguishable groups despite consisting of more than three times as many samples (50 vs 16) and covering a much larger geographical area. Similarly, the western California clade yielded five distinguishable groups after three rounds of fastStructure, while the southwestern California clade yielded two clusters corresponding to the two sampling localities.

**Table 5:** Tajima’s D and nucleotide diversity (π) results. Numbers in parentheses are the number of SNPs considered for each analysis. The number of subgroups refers to the number of groups required to reach the stopping criterion for hierarchical fastStructure analyses.

| Population | Reference length | Tajima's D (SNPs) | $\pi$ (SNPs) | Number of samples | Number of subgroups |
| --- | --- | --- | --- | --- | --- |
| Northeastern California | 1504300 | 0.65 (9296) | 0.0020 (9500) | 11 | 5 |
| Western California | 1545800 | 0.43 (9265) | 0.0016 (9449) | 18 | 4 |
| Eastern California | 1736900 | 0.58 (7718) | 0.0018 (7896) | 5 | 3 |
| Southwestern California | 1760900 | 0.59 (5186) | 0.0010 (5290) | 9 | 2 |
| Northwestern California/Oregon | 782400 | -0.98 (12459) | 0.0022 (12662) | 50 | 3 |
| Combined Samples | 455600 | -1.05 (12938) | 0.0034 (13068) | 93 | NA |

### TreeMix

TreeMix recovered topologies similar to the RAxML tree, with some interesting differences. Model likelihoods improved dramatically until five migration edges were added, and increased modestly thereafter (Figure S2). The best-scoring TreeMix tree with five migration edges and residual errors for each locality are shown in Figure 5, and all other runs are shown in Figures S3-S10 (for locality information, see Table 1). The western California and southwestern California clades always formed reciprocally monophyletic groups, and the long branch between these clades and the remaining samples was used to root all TreeMix topologies. Similarly, the eastern California localities always formed a monophyletic group and were always sister to the northwestern California/Oregon and northeastern California samples.

**Figure 5:**
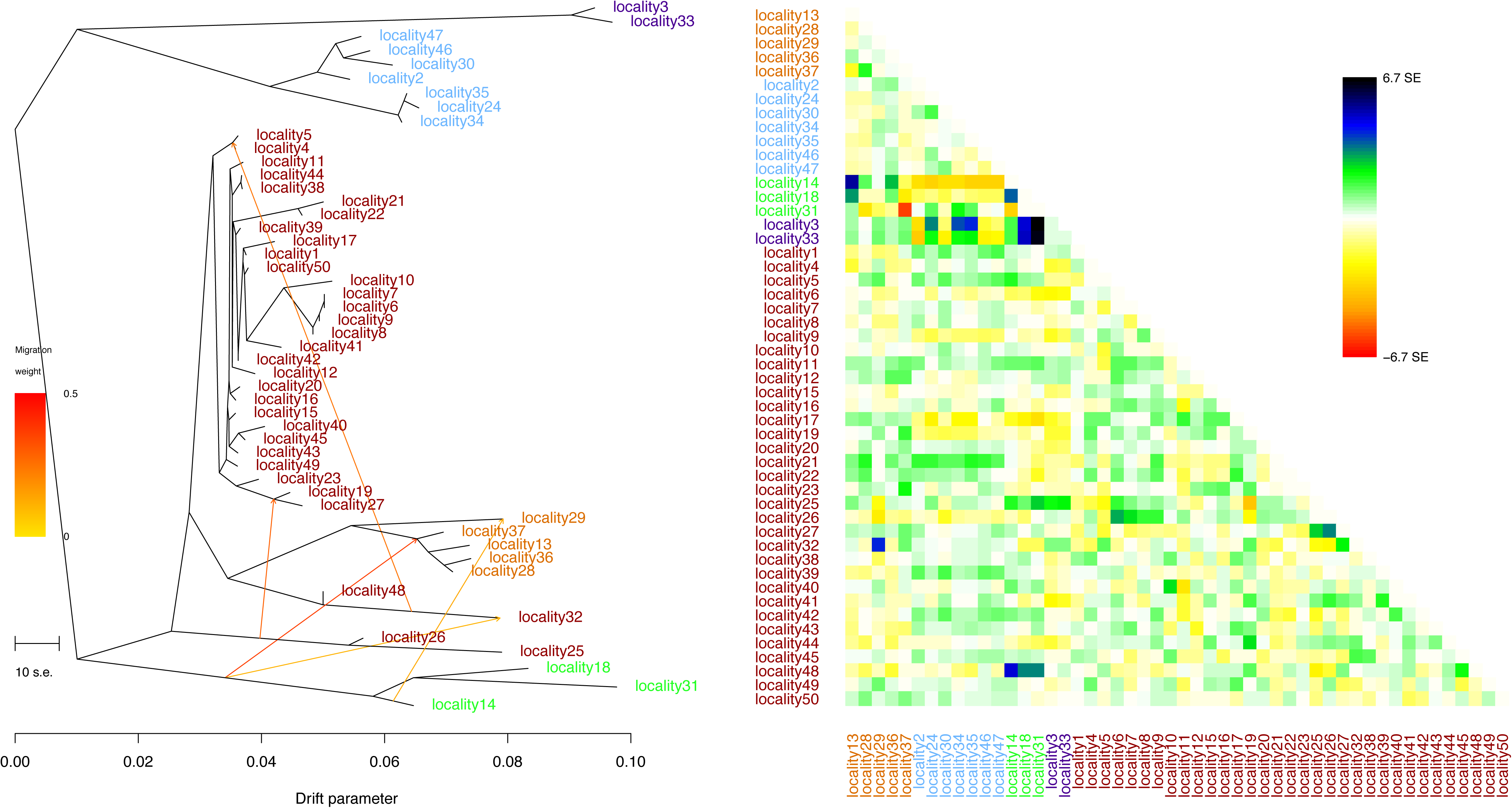
A) TreeMix tree for five modeled migration events. Locality names (Table 1) are colored according to major clade in Figure 1. B) Residuals showing model fit for each locality; localities with small standard errors (SE) fit the TreeMix model better than localities with larger SE values.

The five localities that make up the northeastern California clade always formed a monophyletic group. When no migration was modeled, this group was sister to all northwestern California/Oregon localities (Figure S3), but when at least one migration edge was added the northeastern California localities were nested within the northwestern California/Oregon localities (Figure 5, Figures S4-S10).

The first five added migration edges occurred from the ancestor of the eastern California samples to the ancestor of the northeastern California samples and to the adjacent locality 32 (northwestern California/Oregon), from the ancestor of the central Oregon localities to the southwestern Oregon/far northwestern California localities, and between localities within the northwestern California/Oregon clade (Figure 5). Three-population tests did not indicate that any one of the five major RAXmL clades consist of a mixture of any two of the other four groups (Table S2).

### AMOVA, Fst, and Tajima’s D

Results from AMOVA show that 29.6% of the total genomic variation is partitioned among water basins compared to nearly twice as much (53.9%) among the five major phylogenetic clades (Table 3). Most of the residual variation in the water-basin AMOVA is attributed to variation among localities within water basins (38.0% as opposed to 18.0% in the phylogenetic clade analysis). This suggests that the deep population structure recovered by RAxML, PCA, and fastStructure is not perfectly captured by membership among the different HUC6-level water basins in California and Oregon.

**Table 3:**
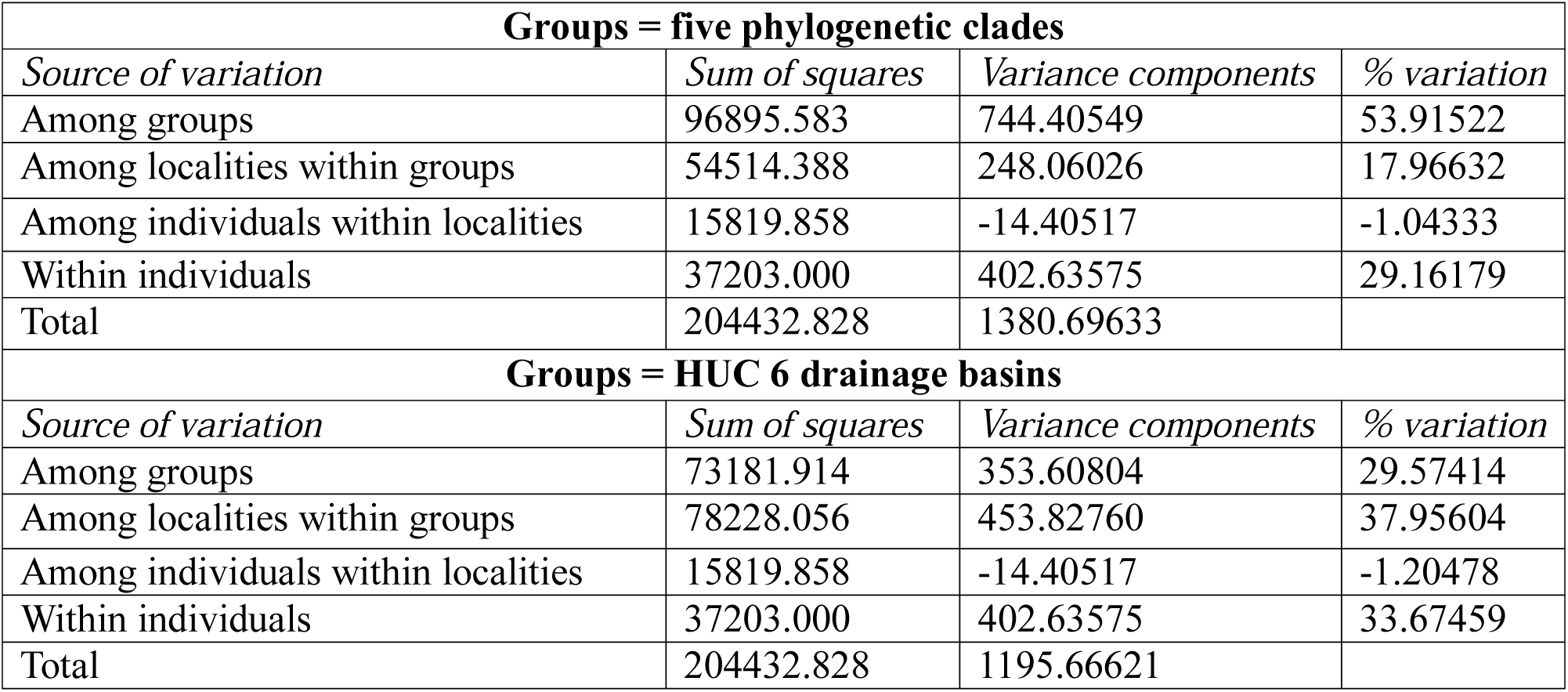
AMOVA results for partitioning sampling localities by phylogenetic clade or by drainage basin.

Fst values were extremely high among the five major clades (Table 4). The lowest Fst was 0.312 for the comparison between northwest California/Oregon and northeast California, and the highest value was 0.794 between the southwestern California samples and the eastern California samples. The southwestern California samples were the most differentiated group from each of the other four groups (average Fst=0.711). Plotting the Fst values of comparisons between localities in the same major clade against geographic distance revealed a strong pattern of genetic isolation by geographic distance within clades (Figure 6A). Fst values were generally at least twice as high for comparisons between localities in different major clades compared to those in the same major clade for the same geographic distance (compare black to colored dots in Figure 6B), and these comparisons tended to be less affected by geographic distance than within-clade comparisons as is expected among highly divergent evolutionary lineages.

**Figure 6:**
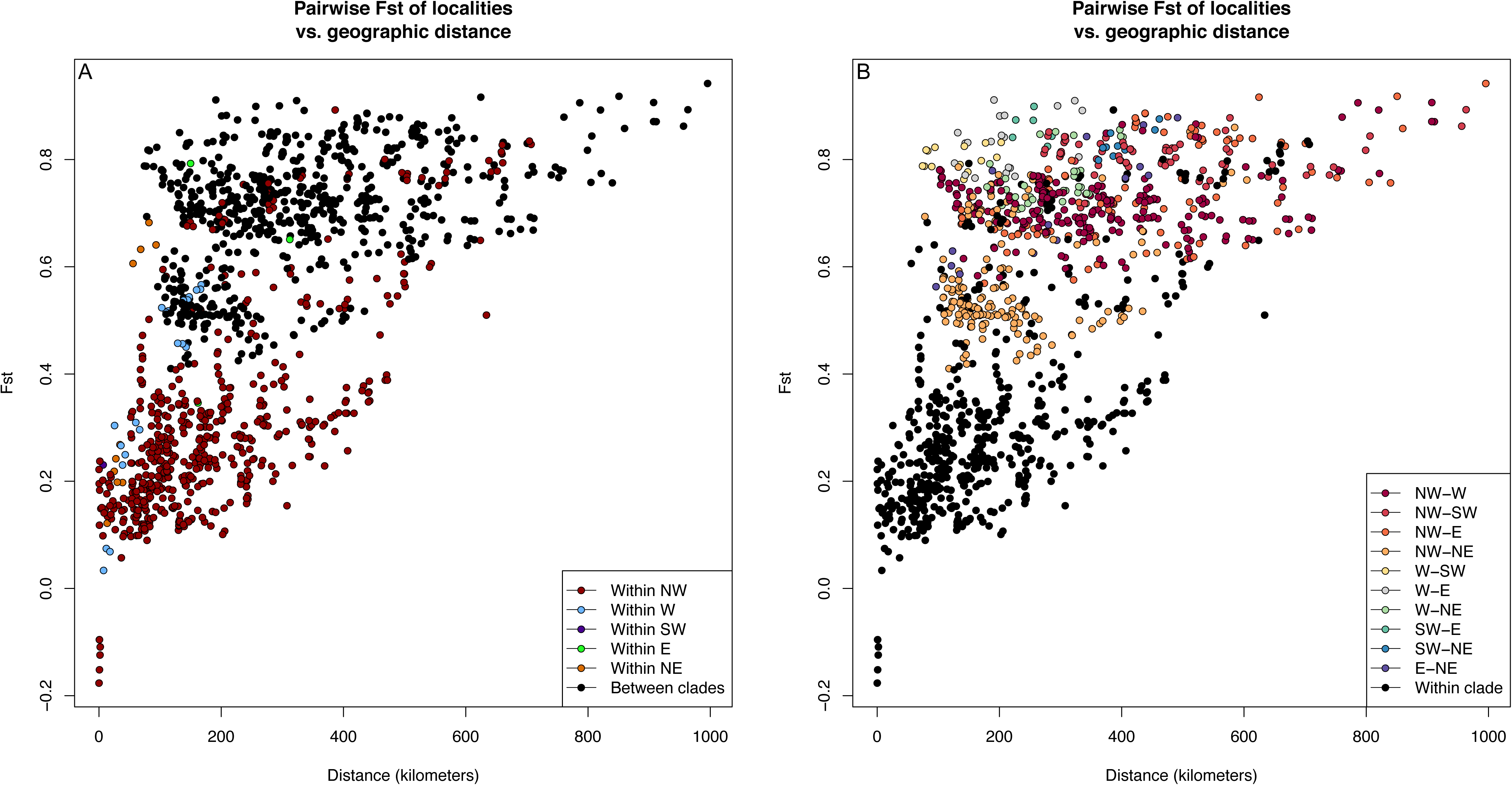
Fst values calculated between sampling localities versus the geographic distances separating localities, coloring dots A) for comparisons within major clades and B) for comparisons between major clades.

**Table 4:** Pairwise Fst values (lower triangle, in bold) and average pairwise JC69 distance values (upper triangle including diagonal, in italics). Population assignment based on the five major clades recovered by RAxML.

|  | Northwest<br>California/Oregon | Eastern<br>California | Western<br>California | Southwestern<br>California | Northeastern<br>California |
| --- | --- | --- | --- | --- | --- |
| Northwest<br>California/Oregon | *0.00113* | *0.00376* | *0.00349* | *0.00460* | *0.00198* |
| Eastern California | **0.503** | *0.00119* | *0.00481* | *0.00562* | *0.00320* |
| Western<br>California | **0.519** | **0.688** | *0.00080* | *0.00417* | *0.00386* |
| Southwestern<br>California | **0.620** | **0.794** | **0.697** | *0.00028* | *0.00509* |
| Northeastern<br>California | **0.312** | **0.545** | **0.630** | **0.734** | *0.00105* |

Tajima’s D values were positive for all clades except for northwestern California/Oregon, which was -0.98 (Table 5), suggesting that this clade may have undergone a recent demographic bottleneck/expansion (Tajima 1989). Nucleotide diversity values ranged from 0.0010 for the southwestern California samples to 0.0034 for northwestern California/Oregon (Table 5). Average sequence distances tended to be greatest when comparing groups to the southwestern California clade, except for the western California clade, which was most distant from the eastern California clade (Table 4).

## Discussion

The central findings of this study were the presence of deep, geographically structured genetic subdivision in *Rana boylii* throughout its range in California and Oregon and the unambiguous biological patterns that were evident in the sequence clustering threshold evaluation metrics. Sequence clustering thresholds exhibited concordant patterns across the four metrics, showing that biological allelic variation began to be split into different clusters at approximately 97% similarity, and thereby supporting 96% as a threshold that splits the maximum number of paralogous regions while minimizing oversplitting.

## Clustering level determination

The choice of clustering threshold is a key parameter in the analysis of RADseq data, and its optimal value depends on the biological system under study. While any RADseq study would benefit from choosing the correct clustering threshold, this parameter is particularly important for organisms with large genomes, which contain a greater number of potentially paralogous regions. Fortunately, the phenomena of oversplitting allelic variants and undersplitting paralogous loci can be evaluated in a RADseq assembly by assessing the relationship of genetic divergence and missingness along a known major axis of genetic variation, as we demonstrate here. Although the metrics that we tested indicate a continuous relationship between each performance criterion and clustering threshold, they also indicate a more precipitous shift in performance at or near 96%. This indicates that most allelic variation in this system is less than 4% divergent, and that 96% is a clustering threshold that serves to maximize the splitting of paralogous loci while minimizing the splitting of allelic variants.

## Phylogeography and population genetics of a deeply structured riverine species

At the coarsest, range-wide level, five deeply divergent genetic groups were present across phylogenetic (Figure 1), ordination (Figure 3) and Bayesian clustering (Figure 4) approaches, even when explicitly modeling admixture (Figure 5). Fst among these groups was extraordinarily high (Table 4), and currently stands as the highest that we are aware of for any anuran (Monson and Blouin, 2004).

The relationships among the five main groups were well resolved with one interesting exception. The northeastern California clade appears most closely related (RAxML with 100% bootstrap support, Figure 1, and TreeMix, Figures 5 and S3-S10) and most similar (using PCA, Figure 3) to the northwestern California/Oregon clade using PCA, but is more closely allied with the eastern California clade in fastStructure analyses. This may hint at a more complex relationship of the northeastern California population with its neighbors to the south and west. Although the three-population test did not indicate that the northeastern California samples are the result of an admixed group between the eastern California samples and those from the northwestern California/Oregon clade (Table S2), TreeMix analyses did recover a signal of migration between ancestors of the eastern California and northeastern California localities (or their ancestors) for all analyses with added migration edges (Figure 5, Figures S4-S10).

At a finer resolution, there are additional indications of deep hierarchical population structure within these five groups. While fastStructure recovered distinct population clusters for each sampling locality in the eastern California and southwestern California clades, the 33 northwestern California/Oregon localities supported just two subunits, one of which spanned more than 50,000 km^2^. The western California localities were intermediate in this regard, with fastStructure recovering five populations across seven localities. Similarly, nucleotide diversity varied by more than a factor of 2 among the major phylogenetic clades, with the western California clade exhibiting a value half that of the northwestern California/Oregon clade. Given the lack of phylogenetic resolution within the northwestern California/Oregon clade and its negative Tajima’s D, both of which suggest a recent population expansion, this high level of nucleotide diversity is consistent with either a range expansion from multiple source populations or some limited admixture from other areas. One concern is that these estimates of genetic diversity may be influenced by sample size (Subramanian 2016) or relative area, as the two southwestern California localities are extremely close together. However, this does not fully explain the depressed π values in the southwestern California samples; the 9 samples from locality 2 (western California clade) and the 6 samples from locality 50 (northwestern California clade) had π values of 0.0012 and 0.0022, respectively (data not shown), which are both higher than the 0.0010 registered for the 9 samples from two southwestern California localities. Therefore, southwestern California samples appear to be undergoing marked reductions in nucleotide diversity compared to other parts of the range (Table 5). Future research could explore this further with demographic genetic modeling approaches.

### Comparison to Lind et al. (2011)

Comparison of our results with a previous, primarily mtDNA study of many of the same localities provides important insights into the increased utility of genome-level SNP data compared to the single locus analyses that characterized earlier phylogeographic and conservation genetic analyses. Both Lind *et al.* (2011) and the current study recovered the genetic distinctiveness of populations at the periphery of the species range in southern Monterey (our southwestern CA) and Kern Counties (part of our eastern California) and in the central Sierra of California (clade C in Lind *et al.* 2011). However, the earlier analyses, based on 1,525 bp of mtDNA, had virtually no power to resolve groupings of populations into more inclusive lineages, and those that were identified has far less resolution and statistical certainty. For example, clade D in Lind *et al.* (2011) is similar to the western California clade identified here, but with modest statistical support (Bayesian posterior probability of 0.95, maximum likelihood bootstrap of 0.67) and included a seemingly inexplicable sample, HBS 37260 from the Sierra foothills. In this study, however, most of these ambiguities are resolved and clarified: the major clades are well-supported, and the substructure that so clearly dominates Sierran and coastal populations south of San Francisco Bay are equally strongly supported. The weak evidence for a recent trans-valley leak, as was recently (and reasonably) suggested by Richmond *et al.* (2014) by a single sample (HBS 37260), can be conclusively interpreted as a case of incomplete lineage sorting of mtDNA, and the confusing placement of samples from northwestern California (del Norte, Humboldt, and Lake Counties) or southwestern Oregon (Curry County) in Lind et al. (2011) resolves unambiguously to Lind *et al.*’s Upper Sacramento River and Marin County/Curry County clade.

One of the major results from Lind *et al.* (2011) was that genetic diversity in *Rana boylii* was largely partitioned by hydrological boundaries, accounting for roughly 40% of the observed genetic variation. While this signal was also present in the current analysis, we recovered a lower proportion of variance described by water basins (30%), while a more informative partitioning by phylogenetic clade explained 54% of the total genetic variation. This may be partially be due to our use of USGS hydrological units as opposed to the hydrological regions employed by Lind *et al.* (2011), but the deep divisions and extraordinary Fst values for our genomic-level data indicate that clade-level historical divergence is the most important component of intraspecific differentiation in *R. boylii*.

## Conservation of an ecological specialist anuran

### Units

Our single strongest recommendation is to manage each of the five major phylogenetic clades (Figure 1) identified herein as independent recovery units of *R. boylii*. The extraordinarily high Fst values among major clades is consistent with the interpretation that *R. boylii* is deeply divided taxon that demands conservation actions following a clade, rather than watershed, approach. Indeed, the among-clade and among-locality Fst values observed here were considerably higher than those found over similar distances in the closely related *Rana cascadae* by Monsen and Blouin (2004), which has previously served as an exemplar of extremely strong genetic differentiation in an amphibian species. While the major hydrological boundaries explain a reasonable portion of the broad-scale genetic diversity of *Rana boylii*, there are also regions where distantly related samples occur in the same HUC6-level hydrological unit. This is particularly true in north-central California, where the northeastern California clade and some northwestern California/Oregon clade localities co-occur. Within these five units, our hierarchical analyses of population structure, visualized in Figure 4, indicate that in four of the five, additional identifiable structure exists down to the level of individual sites. This is true for all clades except the large northwestern California/Oregon group extending from San Francisco Bay north to the California/Oregon border (group 2B, Figure 4). Although our sampling is geographically fairly complete, additional analyses that fill in gaps within and between drainages is necessary to fully identify management subunit boundaries that should help guide management decisions including potential dam removals (Lind et al. 1996, 2016).

### Focus protection and recovery in the southwestern California unit

The southwestern California samples were the most genetically distinct of all the samples analyzed here, driving PC1 and PC2 in Figure 3 despite consisting of just nine samples from two nearby localities. We recovered the lowest nucleotide diversity (π) in the southwestern California samples among the five major clades, and the two sampling localities in this clade separated into two distinct populations in fastStructure analyses despite a distance between sites of only 7km. Contrast this with the bulk of the northwestern California/Oregon samples that formed a single fastStructure population despite covering over 50,000 km^2^. The genetic divergence of these samples emphasizes their extreme importance in conservation and management, given that they likely represent the last vestiges of a population cluster that originally spanned stream habitats from Monterey south to Los Angeles County, a distance of over 500 km (Thomson et al., 2016). The low genetic diversity of the southwestern California samples further points to recent reductions in population size, and emphasizes the challenge they pose for management. Sequencing of formalin-preserved animals collected from this entire block of extirpated *R. boylii* populations is becoming increasingly tractable and should help guide future captive breeding and repatriation efforts (Hykin et al. 2015; Ruane and Austin 2017)

## Acknowledgements

We thank Amy Lind, the Museum of Vertebrate Zoology, University of California, Berkeley, and the Department of Herpetology at the California Academy of Sciences for providing samples, Erin Toffelmier and Genevieve Mount for laboratory assistance, and Phil Spinks for conversations regarding analyses. This work used the Vincent J. Coates Genomics Sequencing Laboratory at UC Berkeley, supported by NIH S10 Instrumentation Grants S10RR029668 and S10RR027303, and the Comet cluster at the San Diego Supercomputing Center--an XSEDE resource (Towns et al. 2014), supported by NSF ACI-1548562. EMM and HBS are supported by NSF-DEB 1257648 and grants from the US Fish and Wildlife Service and the US Bureau of Reclamation. This work was supported by a grant from The Scientific and Technological Council of Turkey (TUBITAK).

## Data Accessibility

Raw sequence reads are deposited at NCBI (PRJNA401430). Assemblies and variant calls from pyRAD are available at doi.org/10.5281/zenodo.885534.

## Author Contributions

EMM analyzed the data and wrote the initial draft manuscript, MG collected the data and edited the manuscript, and HBS developed the initial project and edited the manuscript.

**Figure S1:**
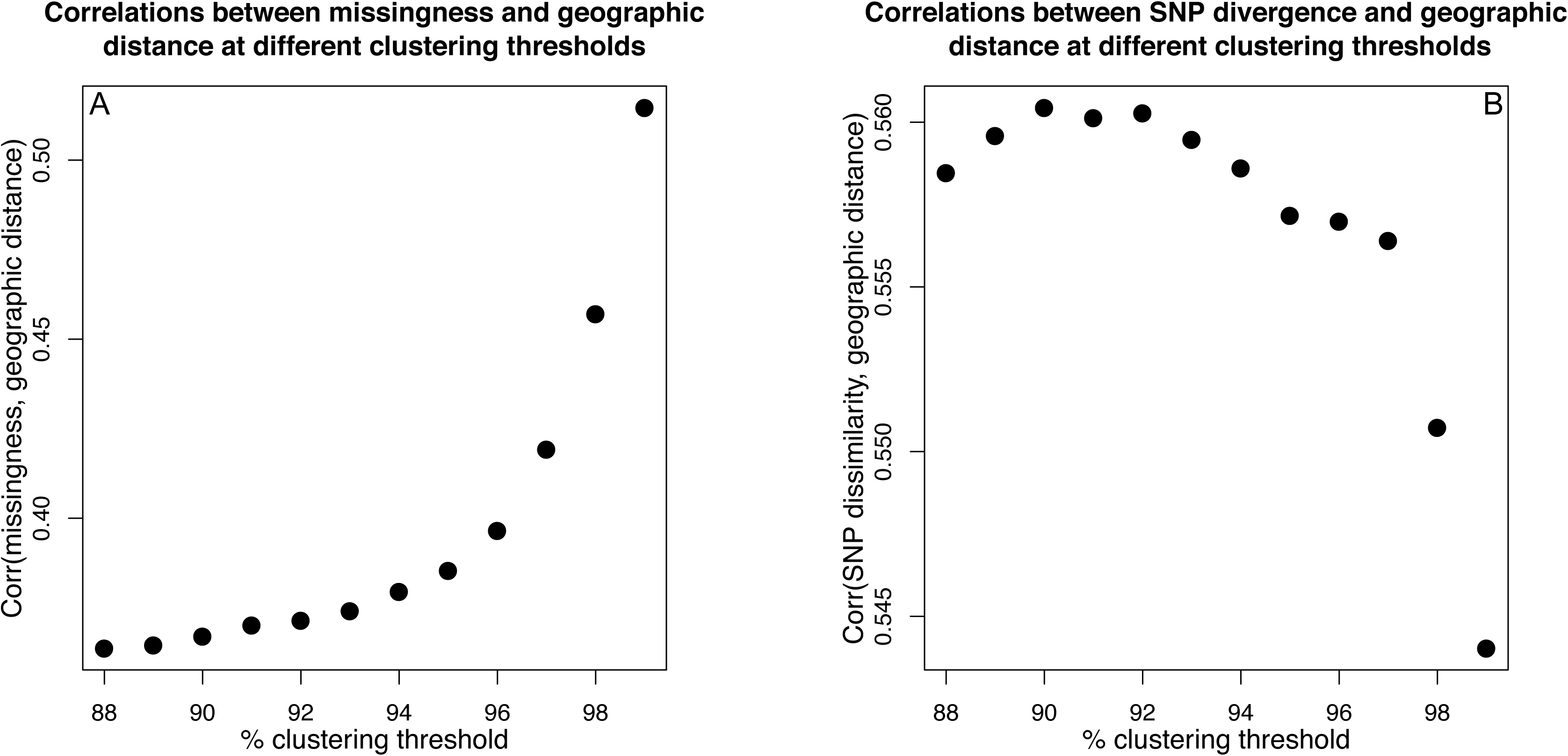
A) Correlation of pairwise data missingness with geographic distance at different pyRAD clustering thresholds. B) Correlation of genetic SNP dissimilarity (1 - identity-by-state of variable sites as calculated by SNPRelate) with geographic distance at different pyRAD clustering thresholds.

**Figure S2:**
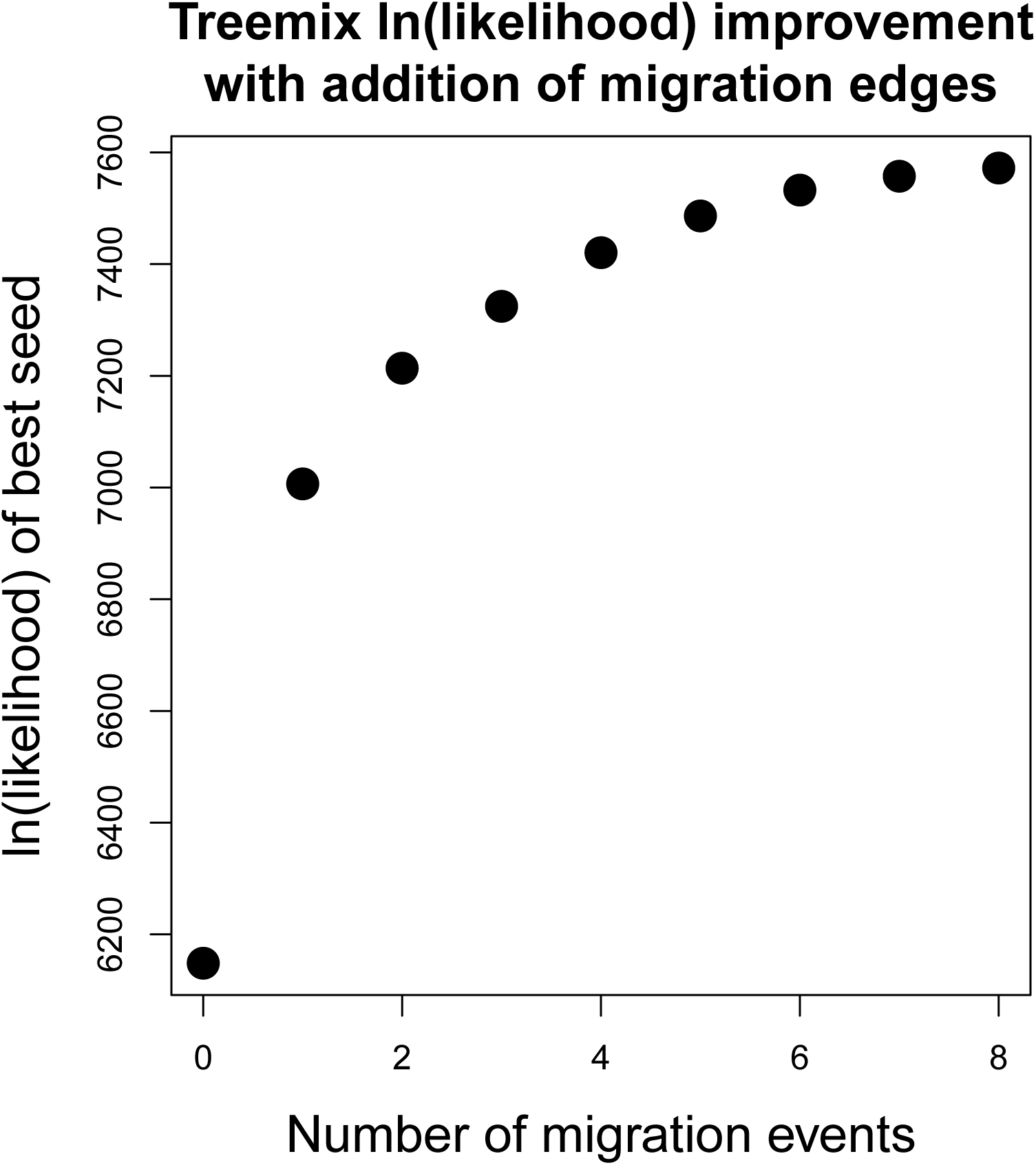
In(likelihoods) of the best of 10 different random number seed runs for the addition of zero to eight added migration edges in TreeMix analyses.

**Figure S3-10:**
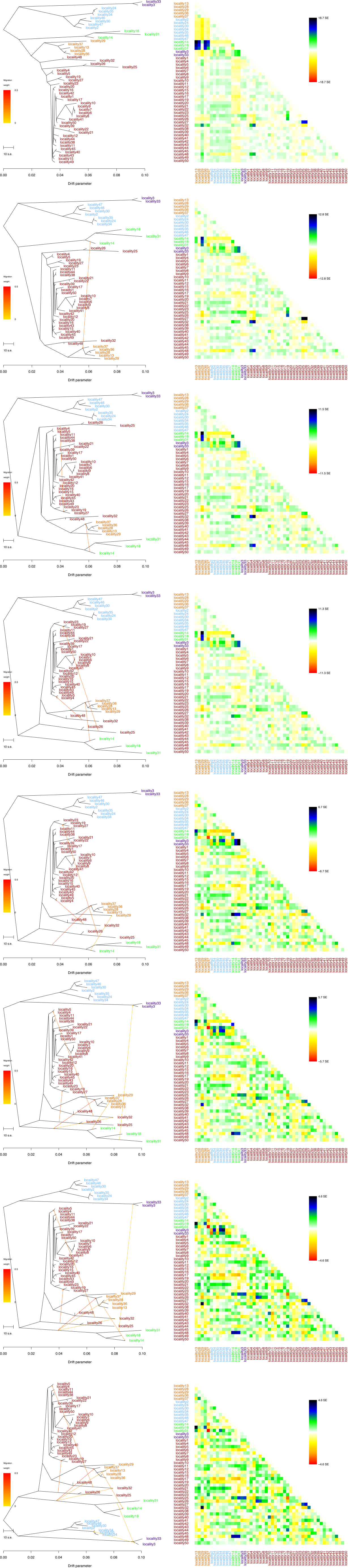
A) TreeMix trees for all modeled migration events except for five (shown in Figure 5). Locality names (Table 1) are colored according to major clade in Figure 1. Locality information is given in Table S1. B) Residuals showing model fit for each locality.

## References

Adams AJ, Kupferberg SJ, Wilber MQ, et al (2017) Extreme drought, host density, sex, and bullfrogs influence fungal pathogen infection in a declining lotic amphibian. Ecosphere. doi: 10.1002/ecs2.1740

Bradford DF, Knapp RA, Sparling DW, et al (2011) Pesticide distributions and population declines of California, USA, alpine frogs, Rana muscosa and Rana sierrae. Environmental Toxicology and Chemistry 30:682–691. doi: 10.1002/etc.425

Danecek P, Auton A, Abecasis G, et al (2011) The variant call format and VCFtools. Bioinformatics 27:2156–2158.

Davidson C (2004) Declining Downwind: Amphibian Population Declines in California and Historical Pesticide Use. Ecological Applications 14:1892–1902. doi: 10.1890/03-5224

Davidson C, Benard MF, Shaffer HB, et al (2007) Effects of Chytrid and Carbaryl Exposure on Survival, Growth and Skin Peptide Defenses in Foothill Yellow-legged Frogs. Environ Sci Technol 41:1771–1776. doi: 10.1021/es0611947

Davidson C, Shaffer HB, Jennings MR (2002) Spatial tests of the pesticide drift, habitat destruction, UV-B, and climate-change hypotheses for California amphibian declines. Conservation Biology 16:1588–1601.

Eaton DAR (2014) PyRAD: assembly of de novo RADseq loci for phylogenetic analyses. Bioinformatics 30:1844–1849. doi: 10.1093/bioinformatics/btu121

Ecoclub Amphibian Group, Pope KL, Wengert GM, et al (2016) Citizen scientists monitor a deadly fungus threatening amphibian communities in northern coastal California, USA. Journal of Wildlife Diseases 52:516–523. doi: 10.7589/2015-10-280

Excoffier L, Lischer HEL (2010) Arlequin suite ver 3.5: a new series of programs to perform population genetics analyses under Linux and Windows. Molecular Ecology Resources 10:564–567. doi: 10.1111/j.1755-0998.2010.02847.x

Excoffier L, Smouse PE, Quattro JM (1992) Analysis of Molecular Variance Inferred from Metric Distances among DNA Haplotypes: Application to Human Mitochondrial DNA Restriction Data. Genetics 131:479–491.

Felsenstein J (1989) PHYLIP-phylogeny inference package (version 3.2). Cladistics 5:163–166. doi: 10.1111/j.1096-0031.1989.tb00562.x

Fu Y-B (2014) Genetic Diversity Analysis of Highly Incomplete SNP Genotype Data with Imputations: An Empirical Assessment. G3 (Bethesda) 4:891–900. doi: 10.1534/g3.114.010942

Gregory TR (2016) Animal Genome Size Database. www.genomesize.com/.

Harvey MG, Judy CD, Seeholzer GF, et al (2015) Similarity thresholds used in DNA sequence assembly from short reads can reduce the comparability of population histories across species. PeerJ. doi: 10.7717/peerj.895

Hayes MP, Jennings MR (1986) Decline of ranid frog species in western North America: are bullfrogs (Rana catesbeiana) responsible? Journal of Herpetology 490–509.

Hoffberg S, Kieran TJ, Catchen JM, et al (2016) Adapterama IV: Sequence Capture of Dual-digest RADseq Libraries with Identifiable Duplicates (RADcap). bioRxiv 44651. doi: 10.1101/044651

Houlahan JE, Findlay CS, Schmidt BR, et al (2000) Quantitative evidence for global amphibian population declines. Nature 404:752–755. doi: 10.1038/35008052

Hykin SM, Bi K, McGuire JA (2015) Fixing Formalin: A Method to Recover Genomic-Scale DNA Sequence Data from Formalin-Fixed Museum Specimens Using High-Throughput Sequencing. PLOS ONE 10:e0141579. doi: 10.1371/journal.pone.0141579

Ilut DC, Nydam ML, Hare MP (2014) Defining Loci in Restriction-Based Reduced Representation Genomic Data from Nonmodel Species: Sources of Bias and Diagnostics for Optimal Clustering. BioMed Research International 2014:e675158. doi: 10.1155/2014/675158

Janes JK, Miller JM, Dupuis JR, et al (in press) The K=2 conundrum. Mol Ecol. doi: 10.1111/mec.14187

Jukes TH, Cantor CR (1969) Evolution of protein molecules. In “Mammalian Protein Metabolism”.(Ed. HN Munro.) pp. 21–132. Academic Press: New York

Kerby JL, Sih A (2015) Effects of carbaryl on species interactions of the foothill yellow legged frog (Rana boylii) and the Pacific treefrog (Pseudacris regilla). Hydrobiologia 746:255– 269.

Kiesecker JM, Blaustein AR, Belden LK (2001) Complex causes of amphibian population declines. Nature 410:681–684.

Kupferberg SJ, Palen WJ, Lind AJ, et al (2012) Effects of Flow Regimes Altered by Dams on Survival, Population Declines, and Range-Wide Losses of California River-Breeding Frogs. Conservation Biology 26:513–524.

Leaché AD, Chavez AS, Jones LN, et al (2015) Phylogenomics of Phrynosomatid Lizards: Conflicting Signals from Sequence Capture versus Restriction Site Associated DNA Sequencing. Genome Biol Evol 7:706–719. doi: 10.1093/gbe/evv026

Lind AJ, Hartwell Jr H, Wilson RPD, others (1996) The effects of a dam on breeding habitat and egg survival of the Foothill Yellow-legged Frog (Rana boylii). Lind AJ, Spinks PQ, Fellers GM, Shaffer HB (2011) Rangewide phylogeography and landscape genetics of the Western US endemic frog Rana boylii (Ranidae): implications for the conservation of frogs and rivers. Conservation Genetics 12:269–284.

Lind AJ, Welsh Jr HH, Wheeler CA (2016) Foothill yellow-legged frog (Rana boylii) oviposition site choice at multiple spatial scales. Journal of Herpetology 50:263–270.

Macey JR, Strasburg JL, Brisson JA, et al (2001) Molecular Phylogenetics of Western North American Frogs of the Rana boylii Species Group. Molecular Phylogenetics and Evolution 19:131–143. doi: 10.1006/mpev.2000.0908

Monsen KJ, Blouin MS (2004) Extreme isolation by distance in a montane frog Rana cascadae. Conservation Genetics 5:827–835. doi: 10.1007/s10592-004-1981-z

Nunziata SO, Lance SL, Scott DE, et al (2017) Genomic data detect corresponding signatures of population size change on an ecological time scale in two salamander species. Mol Ecol 26:1060–1074. doi: 10.1111/mec.13988

Olmo E (1973) Quantitative variations in the nuclear DNA and phylogenesis of the amphibia. Caryologia 26:43–68. doi: 10.1080/00087114.1973.10796525

Peterson BK, Weber JN, Kay EH, et al (2012) Double Digest RADseq: An Inexpensive Method for De Novo SNP Discovery and Genotyping in Model and Non-Model Species. PLoS ONE 7:e37135. doi: 10.1371/journal.pone.0037135

Pickrell JK, Pritchard JK (2012) Inference of Population Splits and Mixtures from Genome-Wide Allele Frequency Data. PLOS Genet 8:e1002967. doi: 10.1371/journal.pgen.1002967

R Core Team (2017) R: A language and environment for statistical computing. R Foundation for Statistical Computing, Vienna, Austria. ISBN 3-900051-07-0, URL www.R-project.org/. www.R-project.org. Accessed 30 Sep 2016

Raj A, Stephens M, Pritchard JK (2014) fastSTRUCTURE: Variational Inference of Population Structure in Large SNP Data Sets. Genetics 197:573–589. doi: 10.1534/genetics.114.164350

Reich D, Thangaraj K, Patterson N, et al (2009) Reconstructing Indian population history. Nature 461:489–494.

Richmond JQ, Backlin AR, Tatarian PJ, et al (2014) Population declines lead to replicate patterns of internal range structure at the tips of the distribution of the California red-legged frog (Rana draytonii). Biological Conservation 172:128–137. doi: 10.1016/j.biocon.2014.02.026

Rodríguez-Ezpeleta N, Bradbury IR, Mendibil I, et al (2016) Population structure of Atlantic mackerel inferred from RAD-seq-derived SNP markers: effects of sequence clustering parameters and hierarchical SNP selection. Mol Ecol Resour 16:991–1001. doi: 10.1111/1755-0998.12518

Rogers SD, Peacock MM (2012) The disappearing northern leopard frog (Lithobates pipiens): conservation genetics and implications for remnant populations in western Nevada. Ecology and Evolution 2:2040–2056.

Roland AB, Santos JC, Carriker BC, et al (2016) Radiation and hybridization of the Little Devil poison frog (Oophaga sylvatica) in Ecuador. bioRxiv 72181. doi: 10.1101/072181

Ruane S, Austin CC (2017) Phylogenomics using formalin-fixed and 100+ year-old intractable natural history specimens. Mol Ecol Resour. doi: 10.1111/1755-0998.12655

Ryan ME, Palen WJ, Adams MJ, Rochefort RM (2014) Amphibians in the climate vise: loss and restoration of resilience of montane wetland ecosystems in the western US. Frontiers in Ecology and the Environment 12:232–240.

Sambrook J, Russell DW (2001) Molecular cloning: a laboratory manual (3-volume set). Cold Spring Harbor Laboratory Press, Cold Spring Harbor, New York

Sexsmith LE (1968) DNA values and karyotypes of Amphibia. PhD thesis. University of Toronto, Ontario

Shaffer HB, Fellers GM, Voss SR, et al (2004) Species boundaries, phylogeography and conservation genetics of the red-legged frog (Rana aurora/draytonii) complex. Mol Ecol 13:2667–2677. doi: 10.1111/j.1365-294X.2004.02285.x

Smith JJ, Putta S, Zhu W, et al (2009) Genic regions of a large salamander genome contain long introns and novel genes. BMC Genomics 10:19. doi: 10.1186/1471-2164-10-19

Stamatakis A (2014) RAxML version 8: a tool for phylogenetic analysis and post-analysis of large phylogenies. Bioinformatics 30:1312–1313. doi: 10.1093/bioinformatics/btu033

Streicher JW, Devitt TJ, Goldberg CS, et al (2014) Diversification and asymmetrical gene flow across time and space: Lineage sorting and hybridization in polytypic barking frogs. Mol Ecol 23:3273–3291. doi: 10.1111/mec.12814

Stuart SN, Chanson JS, Cox NA, et al (2004) Status and trends of amphibian declines and extinctions worldwide. Science 306:1783–1786. doi: 10.1126/science.1103538

Subramanian S (2016) The effects of sample size on population genomic analyses – implications for the tests of neutrality. BMC Genomics. doi: 10.1186/s12864-016-2441-8

Tajima F (1989) Statistical method for testing the neutral mutation hypothesis by DNA polymorphism. Genetics 123:585–595.

Thomson RC, Wright AN, Shaffer HB (2016) California Amphibian and Reptile Species of Special Concern. University of California Press, Oakland, California

Towns J, Cockerill T, Dahan M, et al (2014) XSEDE: Accelerating Scientific Discovery. Computing in Science & Engineering 16:62–74. doi: 10.1109/MCSE.2014.80

US Fish and Wildlife Service (2015) Endangered and Threatened Wildlife and Plants; 90-Day Findings on 31 Petitions. 80 FR 37568 37568–37579.

USDA-NRCS, USGS, EPA (2016) The Watershed Boundary Dataset (WBD) was created from a variety of sources from each state and aggregated into a standard national layer for use in strategic planning and accountability. Watershed Boundary Dataset for the United States of America. Available URL:<ftp://rockyftp.cr.usgs.gov/vdelivery/Datasets/Staged/Hydrography/WBD/National/GDB/>.

Vähä J-P, Erkinaro J, Niemelä E, Primmer CR (2007) Life-history and habitat features influence the within-river genetic structure of Atlantic salmon. Molecular Ecology 16:2638–2654. doi: 10.1111/j.1365-294X.2007.03329.x

Vinogradov AE (1998) Genome size and GC-percent in vertebrates as determined by flow cytometry: The triangular relationship. Cytometry 31:100–109.

Vredenburg VT, Bingham R, Knapp R, et al (2007) Concordant molecular and phenotypic data delineate new taxonomy and conservation priorities for the endangered mountain yellow-legged frog. Journal of Zoology 271:361–374.

Weir BS, Cockerham CC (1984) Estimating F-statistics for the analysis of population structure. evolution 1358–1370.

Whiles MR, Hall RO, Dodds WK, et al (2013) Disease-driven amphibian declines alter ecosystem processes in a tropical stream. Ecosystems 16:146–157.

Zheng X, Levine D, Shen J, et al (2012) A high-performance computing toolset for relatedness and principal component analysis of SNP data. Bioinformatics 28:3326–3328. doi: 10.1093/bioinformatics/bts606

